# A 128-channel receive array with enhanced SNR performance for 10.5 tesla brain imaging

**DOI:** 10.1101/2024.10.20.619294

**Authors:** Russell L Lagore, Alireza Sadeghi-Tarakameh, Andrea Grant, Matt Waks, Edward Auerbach, Steve Jungst, Lance DelaBarre, Steen Moeller, Yigitcan Eryaman, Riccardo Lattanzi, Ilias Giannakopoulos, Luca Vizioli, Essa Yacoub, Simon Schmidt, Gregory J. Metzger, Xiaoping Wu, Gregor Adriany, Kamil Ugurbil

**Affiliations:** Center for Magnetic Resonance Research (CMRR), University of Minnesota, Minneapolis, MN 55455; Center for Advanced Imaging Innovation and Research (CAI2R) and Bernard and Irene Schwartz Center for Biomedical Imaging, Department of Radiology, New York University Grossman School of Medicine, New York, NY, USA

**Keywords:** ultra-high field MRI, UHF, receive array, 128-channel, RF coil, neuroimaging

## Abstract

**Purpose:** To develop and characterize the performance of a 128-channel head array for brain imaging at 10.5 tesla and evaluate the potential of brain imaging at this unique, >10 tesla magnetic field.

**Methods:** The coil is composed of a 16-channel self-decoupled loop transmit/receive array with a 112-loop receive-only (Rx) insert. Interactions between the outer transmitter and the inner 112Rx insert were mitigated using coaxial cable traps placed every 1/16 of a wavelength on each feed cable, locating most preamplifier boards outside the transmitter field and miniaturizing those placed directly on individual coils.

**Results:** The 128-channel array described herein achieved 77% of ultimate intrinsic SNR in the center of the brain. Transmit field maps obtained experimentally on a phantom with and without the receive array were similar and matched EM simulations, leading to FDA approval for human imaging. Anatomical and functional data, including with power demanding sequences, were acquired successfully on human volunteers.

**Conclusions:** Counterintuitive to expectations based on magnetic fields ≤7T, the higher channel counts provided SNR gains centrally, capturing ∼80% uiSNR. Fraction of uiSNR achieved centrally in 64Rx, 80Rx, and 128Rx arrays suggested that a plateau was being reached at 80%. At this plateau, linear to approximately quadratic B_0_ dependent SNR gains for the periphery and the center, respectively, were observed for 10.5T relative 7T.

## 1. INTRODUCTION

The introduction and subsequent rapid evolution of MRI has revolutionized clinical diagnosis and biomedical research in humans, particularly for the human brain. In this effort, ultrahigh magnetic fields (UHFs, defined as ≥7T) increasingly play a critical role due to the presence of magnetic field dependent gains in signal-to-noise ratio (SNR) and, in some cases, contrast-to-noise ratio (CNR). These gains have proven to be particularly important for functional magnetic resonance imaging (fMRI) (e.g.^1–7^), and increasingly for imaging brain anatomy (e.g.^8–13^), connectivity,^14,2^ and pathologies (e.g. ^15–18^). These accomplishments have motivated the development of >10T systems for human imaging, exemplified by the 10.5T effort in our laboratory,^19–24,13,25^ the 11.7T system in France,^26,27^ and the recent 14T initiative in Netherlands.^28^

The SNR gains at UHFs are best understood in the context of ultimate intrinsic SNR (uiSNR)^29–34^ which is the theoretically maximal SNR within a particular sample at a given magnetic field, independent of any RF coil specifics. Capturing the uiSNR in the human head at UHFs generally relies on multichannel receiver arrays composed of loops tiled over a close-fitting head-conformal former.^35–46^ EM simulations of an ideal coil geometry,^47^ as well as experimental data at 3T^35^ and 7T^41,45^ indicate that thirty-two loops would capture most of the central uiSNR in the human head. In contrast, recent EM modeling and experimental data demonstrated that 32- or 64-channel receive (Rx) arrays composed of loops capture a smaller fraction of the central uiSNR at 10.5T compared to 7T,^48,13,25^ and that an 80Rx array^13^ showed significant increases in central SNR relative to the 64Rx array alone, while still only capturing ∼71% of the central uiSNR. These results suggest that further significant gains centrally may be possible with higher channel counts, consistent with the expectation that a greater number of smaller receive elements would better approximate the ideal current patterns necessary to approach the uiSNR^49^ at very high magnetic fields. There are additional reasons for pursuing a high channel-count head arrays at UHFs. These include potential gains in peripheral SNR^47,41,45^ and parallel imaging through suppression of the g-factor noise (e.g. ^41,45^).

In this paper we describe the construction and performance of a 128-channel receive 16-channel transmit (128Rx/16Tx) array operating at 447 MHz, the ^1^H Larmor frequency at 10.5T. Such an array faces major challenges including the practical problems of manufacturing the complex structure and managing the coupling between the plethora of cables and other conductors in the receive array with the circumscribing transmitter. The latter can suppress the delivery of sufficient power to the sample, damage receiver electronics, compromise the performance of the receive elements and, most importantly, preclude the generation of accurate EM models of the transmitter, a necessary step in defining safe operational limits in human experiments. We present solutions to these challenges, leading to a 10.5T 128Rx/16Tx array approved by the FDA for human imaging, and first time human anatomical and functional images acquired by such a high channel count array at this magnetic field.

## 2. METHODS

The 128Rx/16Tx array described in this paper is composed of an inner 112-channel receive only (112Rx) array and a circumscribing 16-channel transceiver (16Tx/Rx). It largely follows the mechanical design concept of previous 32Rx or 64Rx arrays.^50,51,13^ However, both the 16Tx/Rx and the 112Rx arrays also depart in many ways from previous works as described below.

### 2.1 Receiver Array

The 112Rx array (Figure 1) was constructed onto a PETG plastic head-conformal former designed in SolidWorks (Dassault Systèmes, Vélizy-Villacoublay, FR) and fabricated with an F410 (Fusion 3, Greensboro, NC, USA) fused deposition modeling printer; it approximated the surface of our previous coil formers^13^ but was larger, especially over the frontal areas (Figure S1). The surfaces, however, had zero gaussian curvature which ensured that sheets of material can be wrapped onto them without distortion or wrinkling. This permitted loop conductors designed in 2D CAD (Autodesk Eagle, San Francisco, CA, USA) and fabricated onto flexible printed circuit (FPC) laminates that could then be wrapped onto the former (Figure S2). The 3D model is available in the Supporting Information.

**Figure 1:**
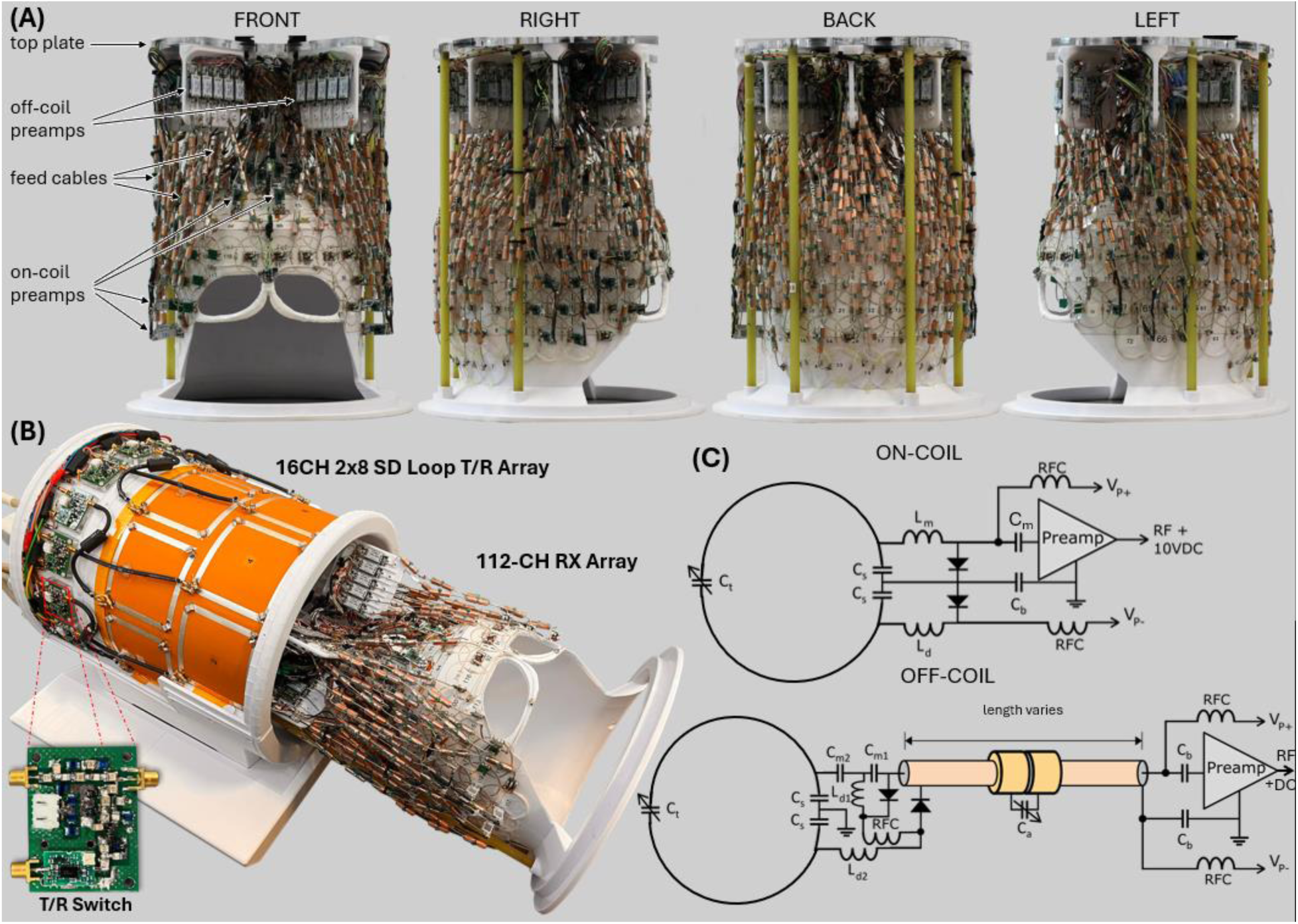
(A) Shows the completed 112-channel receive-only array insert from four different angles to reveal different construction techniques employed in each region of the array with key points of interest labelled. (B) This 112Rx array is shown with its 16-channel self-decoupled loop transceiver with integrated T/R switches allowing for 16 T/R channels. Receive array is partially removed for display purposes only. (C) Schematics for the on-coil and off-coil preamp receive channels.

The 112 loops were arranged in 7 rows: 99 loops on FPCs and 13 loops constructed from 1.0 mm diameter silver-plated copper wire. Eleven wire loops complete the row on top of the head while two are large eye loops. The lower four rows use 40 mm diameter round loops of two segments with overlap nearest neighbor decoupling.^52^ The top three rows are of varying sizes and shapes including oval loops, square loops, and wedge/triangular loops. These have four segments.

A plastic pillar extends from the former upwards 14 cm to a 10mm-thick polycarbonate top plate. There are also four ∼9.5 mm diameter G-10 fiberglass rods that act as struts between the bottom ring of the former and the top plate (Figures 1 and S2). The top plate provides mechanical support for the preamp tree, bias tee mounts, and four strain relief glands for coil cable bundles. Four system plugs are each populated with 32Rx coaxial connections, 16 PIN diode DC connections, and 3 coil code pins (ODU, Mühldorf, DE).

There are 57 on-coil and 55 off-coil preamps. The 112Rx and transceiver formers are close fitting in the posterior and anterior aspect, so off-coil preamps (mounted to the preamp tree) were employed for these loops, while on-coil preamps were used where space allowed, primarily on the left, right, and top of the former.

The preamp used is the WMA447D (WanTcom, Chanhassen, MN, USA) with a gain of 28 dB, NF of 0.5 dB, and Z_in_=1.5 ohm. Low-loss, semi-rigid 1.2 mm diameter coaxial cables (UT47C-LL-TP, AmphenolCIT, St. Augustine, FL, USA) were used for all channels below the top-most row. Interconnects were provided on the preamp tree to convert from the 2mm flexible receive coax (Leoni AG, Nuremberg, DE) to the semi-rigid coax.

For channels employing off-coil preamps, all loop feed circuits (Figure 1C) use a pair of equal-value capacitors (*C*_s_) to produce a virtual ground point to which the feed coaxial cable ground is connected. Matching is achieved with series capacitance (C_m1_ & C_m2_). Posterior-located loops use a miniature 5x5x0.8mm^3^ (WxLxThk) feed PCB populated with 0603 SMD capacitors (Knowles- Syfer, Itasca, IL, USA), 0603 ceramic RF choke (0603CS series, CoilCraft Inc., Cary, IL, USA), 0807 SMD square air core detune inductor (CoilCraft 0807SQ series), and a single PIN diode (MA4P1250NM-1072T, MACOM, Lowell, MA, USA) for detune. The anterior half of the coil has larger feed circuit PCBs that accommodate 1111 SMD capacitors (100B series, American Technical Ceramics, Huntington Station, NY, USA), 1008 SMD RF choke (1008CS series, CoilCraft), two detune inductors (0806SQ/0807SQ/0908SQ series, CoilCraft) and two PIN diodes. This larger feed board both detunes the loop (L_d2_||C_s_) and offers an LC resonant block (L_d1_||C_m1_) for preamp protection during the transmit pulse.

On-coil preamp feed circuitry (Figure 1C) also uses a pair of capacitors (C_s_) to create a virtual ground on loop to which the preamp ground can be referenced. Impedance matching and phase transformation are accomplished with inductor L_m_, the PIN diode reverse bias capacitance, and capacitor C_m_ between the loop and preamp input. During transmit, two PIN diodes provide receiver protection: one detunes the loop and another clamps the RF input at the preamp. The on-coil preamp board also implements bias tees for the PIN diode biasing.

Off-coil preamps use a quarter wavelength transformation for preamp decoupling.^52^ Because the preamp tree is further from the loop conductors than one quarter wavelength, but closer than three-quarters wavelength, lumped-component low-pass pi-network phase shifters are used to achieve the nearest multiple of n*λ/4 (where n is an odd integer) and ensure that a high impedance is presented at the coil.

A critical design goal was to minimize the influence of the 112Rx array on the transmit field. Any conductor within the transmit field with a length equivalent to a significant fraction of a wavelength (i.e., >λ/10) could re-radiate currents induced by the transmitter. Two major sources of such interaction were identified: the output cabling (after the preamp) and the feed cabling to the receive loops (between the coil and preamp). The sheaths of output cabling which branches in many directions from a common origin (the cable bundle) appears like an end cap RF mirror in the same way as ground radials serve as an RF mirror for large navigational monopole antennas. The output cabling and off-coil preamps were positioned well outside the volume of the transmit loops. The consequence of this decision was longer feed cables. These conductors, both DC wiring as well as coaxial cables, were segmented using high impedance circuitry every λ/16 (∼42 mm). RF chokes were used on PIN diode DC lines to produce this high impedance, while coaxial cable traps were implemented on coaxial feed cables. A single variable capacitor was used to tune each cable trap.^53^ Individual RF coil elements in the array could also interact with the transmit field; however, they are already short relative to the wavelength to ensure uniform current distribution.

### 2.2 Transceiver Array

RF transmission was accomplished with a 16Tx/Rx transceiver set up to transmit and receive with each of its elements. This array is based on self-decoupled (SD) loops,^54^ and was described previously.^13^ For convenience, we also include a detailed description as Supporting Information.

### 2.3 EM Simulation and Safety Validation

The 10.5T scanner operates under an Investigational Device Exemption (IDE) from the FDA, requiring each RF coil to be FDA approved before use with humans. The EM simulation and safety validation necessary for this process was described in detail in our previous work^13,25^ for an identical transmitter structure (also see Supporting Information).

### 2.4 Data Acquisition

Coil characterization data and images were obtained on a MAGNETOM 10.5T (Siemens Healthineers, Erlangen, DE) system based on an 88 cm warm-bore magnet (Agilent Technologies, Oxford, UK), configured with a Siemens SC72D body gradient (70 mT/m maximum amplitude, 200 T/m/s slew rate). The transmit RF is supplied from a 16-channel, 2 kW/channel power amplifier (Stolberg HF-Technik AG, Stolberg, DE). The console is equipped with a 128- channel receiver developed in-house with Siemens-supplied components. The patient table was modified in-house to accommodate three 32-channel system plugs in addition to the standard system plugs.

Coil characterization used a lightbulb-shaped phantom mimicking the dimensions and electrical properties of the human head (σ=0.65 S/m, Ɛ_r_=47.2); it was described in detail in our previous work.^13,25^

### 2.5 Benchtop Measurement

Coil elements were tuned and characterized on the bench using a 16-port vector network analyzer (ZNBT8, Rohde & Schwarz, Munich, DE). Coils were probed using in-house fabricated single and decoupled double B-field probes. Quality factors were evaluated based on a transmission-based (S_21_) 3 dB bandwidth method. The completed coil system was inspected for spurious emissions (indicative of LNA oscillations) using a spectrum analyzer (FieldFox N9914A, Keysight, Santa Rosa, CA, USA)

### 2.6 SNR Measurement and g-Factor Calculations

Channel-wise relative B_1_^+^ maps were acquired in vivo using the technique described by Van de Moortele et al.^55^ Subsequently, phase-magnitude B_1_^+^ shimming was performed following the nonlocalized efficiency cost-function of He et al.^56^ targeting the entire cerebrum. Absolute B_1_^+^ maps were acquired using actual flip-angle imaging.^57^ SNR maps were calculated from gradient echo images (TE/TR=3.8/10000 ms, FA=90°, FOV=256x208 mm^2^, voxel size=2x1x2 mm³) in combination with identical noise images acquired without excitation voltage, consistent with previous techniques.^13^ Array performance was evaluated using the lightbulb-shaped phantom by comparing experimentally measured SNR against the uiSNR.^29–34^ The uiSNR was calculated for the lightbulb-shaped phantom as previously described.^13,25^

g-factor maps were calculated from the SENSE equation^58^ for simultaneous multi-slice (SMS)/multiband (MB) acquisition^59–62^ with slice and phase-encode undersampling, as described previously.^41^ g-factor data were obtained in vivo with a fully sampled 1-mm isotropic 3D GRE sequence, and ESPIRIT^63^ was used for sensitivity profile calculation. For visualization, maximum intensity projections (MIP) of g-factors were extracted in the left-right (LR) direction over an 80- mm thick slab in the sagittal plane and displayed over a silhouette of the central sagittal slice as 1/g maps.

### 2.7 Anatomical Imaging

Susceptibility-weighted images (SWI) were acquired in 36 contiguous axial-coronal oblique slices with 0.2x0.2 mm^2^ in-plane and 1.3 mm slice resolution using 3D gradient recalled echo (GRE) (FOV=205x188x46.8 mm³; iPAT=5; TE/TR=18/35 ms; TA∼300 s).

Anatomical T_2_*-weighted images were acquired with the GRE method at 0.2 mm in-plane and 1 mm slice resolution. Two partial brain volumes were acquired in both sagittal (FOV=214x214x30 mm³) and axial (FOV=178x204x28.5 mm³) slice orientations (TE/TR=15.5/600 ms; FA=35°; iPAT=2; TA=288 s).

### 2.8 Functional Imaging

Functional imaging of brain activity (fMRI) was obtained with 3D-GRE echo-planar imaging (EPI) imaging developed in CMRR and publicly is available (http://www.cmrr.umn.edu/multiband/). Acquisitions employed single shot to cover k_x_-k_y_ space with four-fold undersampling in the phase encode dimension while the third dimension (k_z_) was fully segmented. Whole brain 3D-GRE-EPI images obtained with 0.8 mm isotropic resolution (FOV=187x200x112 mm^3^; iPAT=4; TE/TR=20/83 ms; Volume Acquisition time (VAT)∼12s) are presented to demonstrate EPI quality over the entire brain; fMRI data with visual stimuli were acquired with the same 3D-GRE-EPI sequence but over an axial slab with 0.8 mm isotropic resolution; FOV=148x200x35 mm^3^; iPAT=4; TE/TR=17/74 ms; VAT∼3.5 s.

## 3. RESULTS

### 3.1 Benchtop measurements

Figure 2 displays ratio of unloaded to loaded Q (Q-ratio=Q_U_/Q_L_) for loops tuned to proton Larmor frequency for 7 and 10.5T (298 and 447MHz, respectively) as a function of loop diameter and distance from the sample. The diameter of the loops in the 128Rx array is ∼4 cm, which appears well loaded by the sample (Figure 2A). However, the distance to the sample has a significant impact, indicating that not all the elements of the inner 112Rx array will be equally efficient. Q-ratios rapidly increase with coil diameter at both magnetic fields and are higher for 10.5T for all diameters measured; the difference gets larger with closer proximity to the sample (Figure 2B).

**Figure 2:**
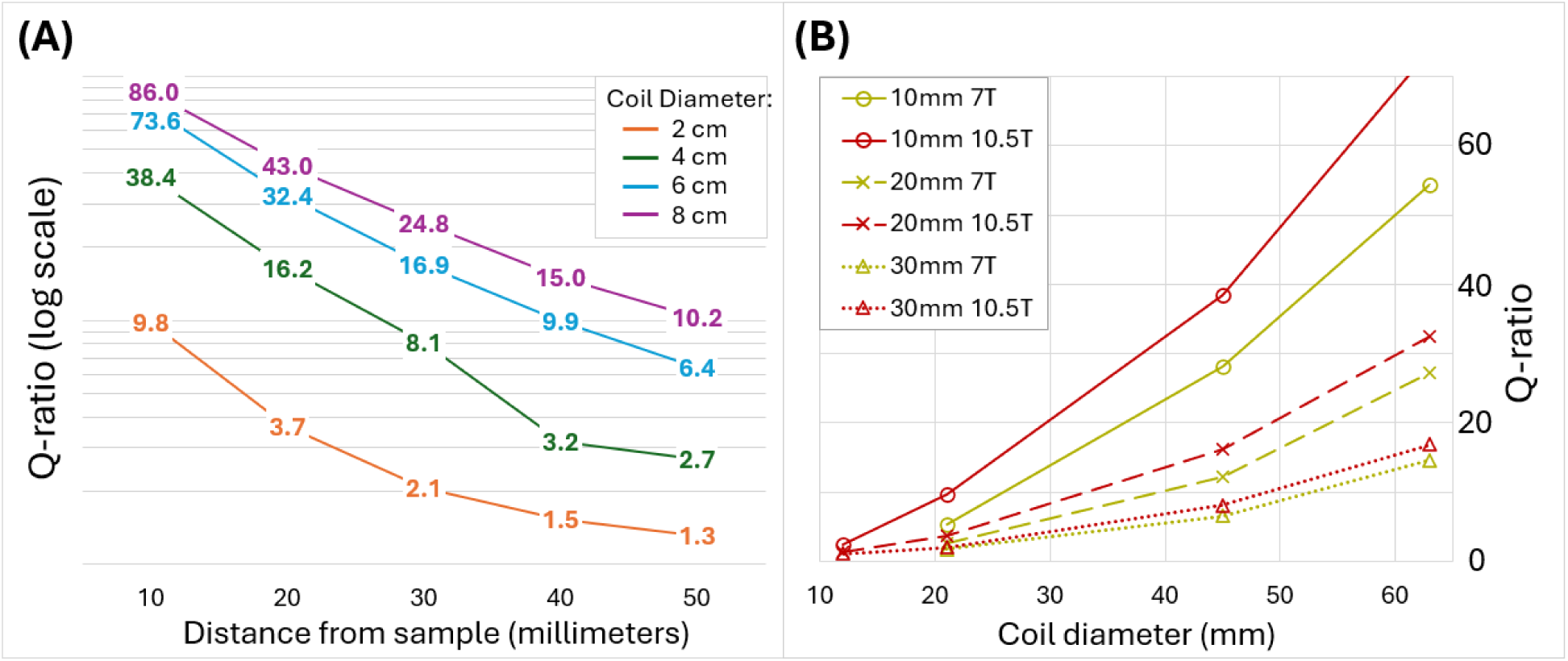
Q-ratio (ratio of unloaded to loaded Q (Q*_U_*/Q*_L_*)): (A) at 10.5T for four different diameter loops (2, 4, 6, and 8 cm) as a function of distance from the loading sample (note the log scale on the vertical axis), and (B) as a function of coil diameter at 7T (yellow) and 10.5T (red) for three different separations from the sample: 10 mm (solid line), 20 mm (dashed line), and 30 mm (dotted line).

Additional benchtop measurements included S-parameters. Simulated S-parameters for the 16Tx/Rx transceiver (without the 112Rx) are shown together with the measured S-parameters (with the 112Rx inserted) in Figure 3, demonstrating good agreement. A full S-parameter matrix was not measured for the 112Rx array. Instead, typical performance metrics were measured for several channels. Q_U_ for the 4 cm diameter round flexible printed circuit loop ranged from 100 to 120 while typical Q_L_ ranged from 40 to 60, yielding Q-ratios ranging from below 2.0 for loops furthest from the load, to about 3.0 for loops closest to the load.

**Figure 3:**
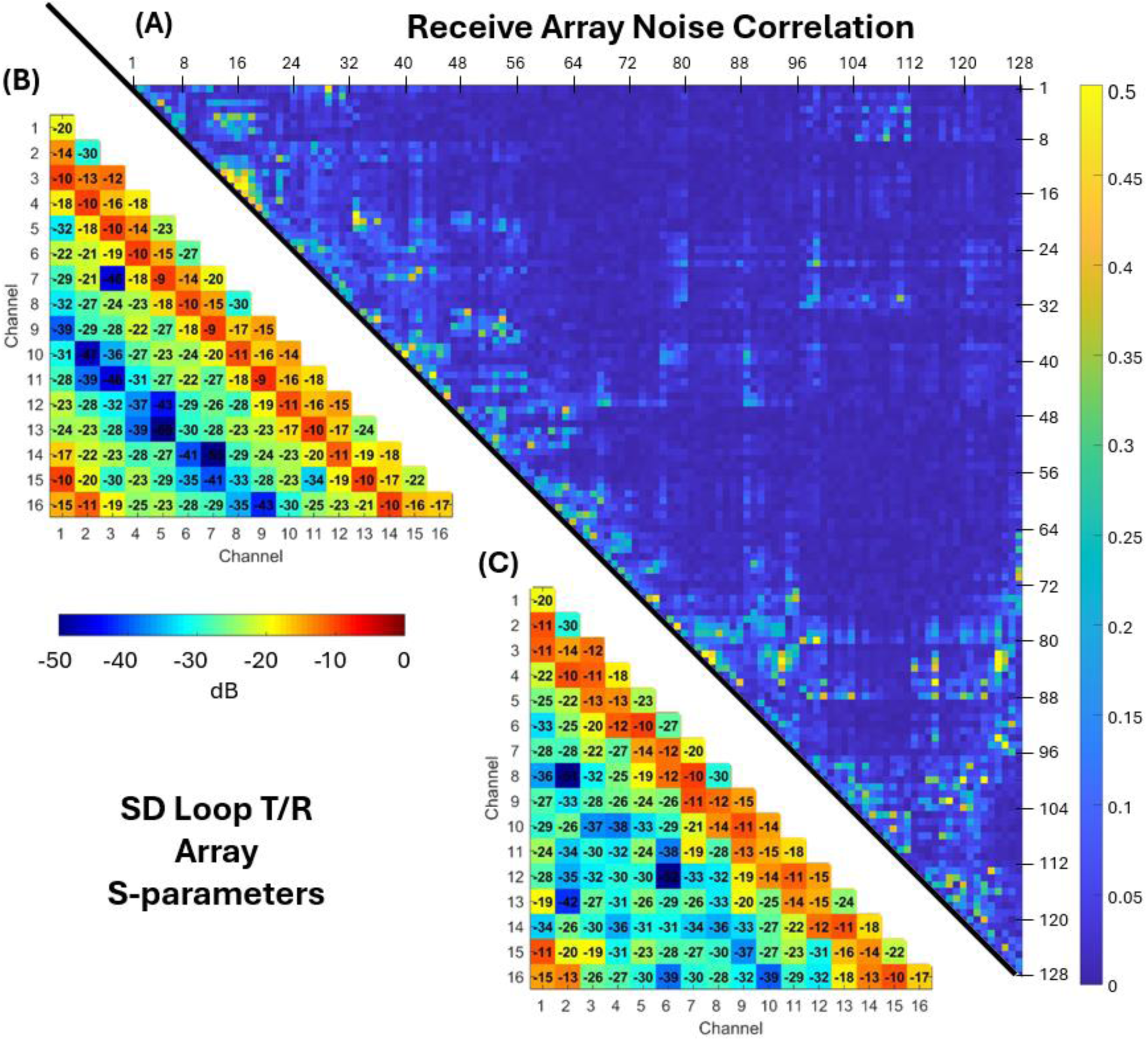
(A) Noise correlation matrix for all channels of the 128-channel receive array (note the scale only goes up to 0.5). (B) Simulated and (C) bench measured S-parameters (dB magnitude scale) of the 16-channel T/R array.

Receive loops were matched to a return loss < -12 dB. Without preamp decoupling, typical nearest neighbor coupling was -15 dB, while next-nearest neighbor coupling was ∼-6 dB. Preamp decoupling reduced coupling a further 16 dB. Loops detuned by 20 dB for a single PIN diode and >40 dB for two PIN diodes. Insertion losses between loop and preamp range from <0.1 dB up to 0.4 dB, dependent upon the precision of tuning on lumped-component phase-shifters.

### 3.2 Interactions between theTransceiver and Receive-only Arrays

Transceiver and receiver array interactions were evaluated using maps of transmit efficiency 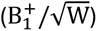 with and without the 112Rx array placed within the transceiver array (Figure 4). The B_1_^+^ maps provided include a circularly polarized (CP) shim configuration accompanied by two pseudo-random phase-only shims.

**Figure 4:**
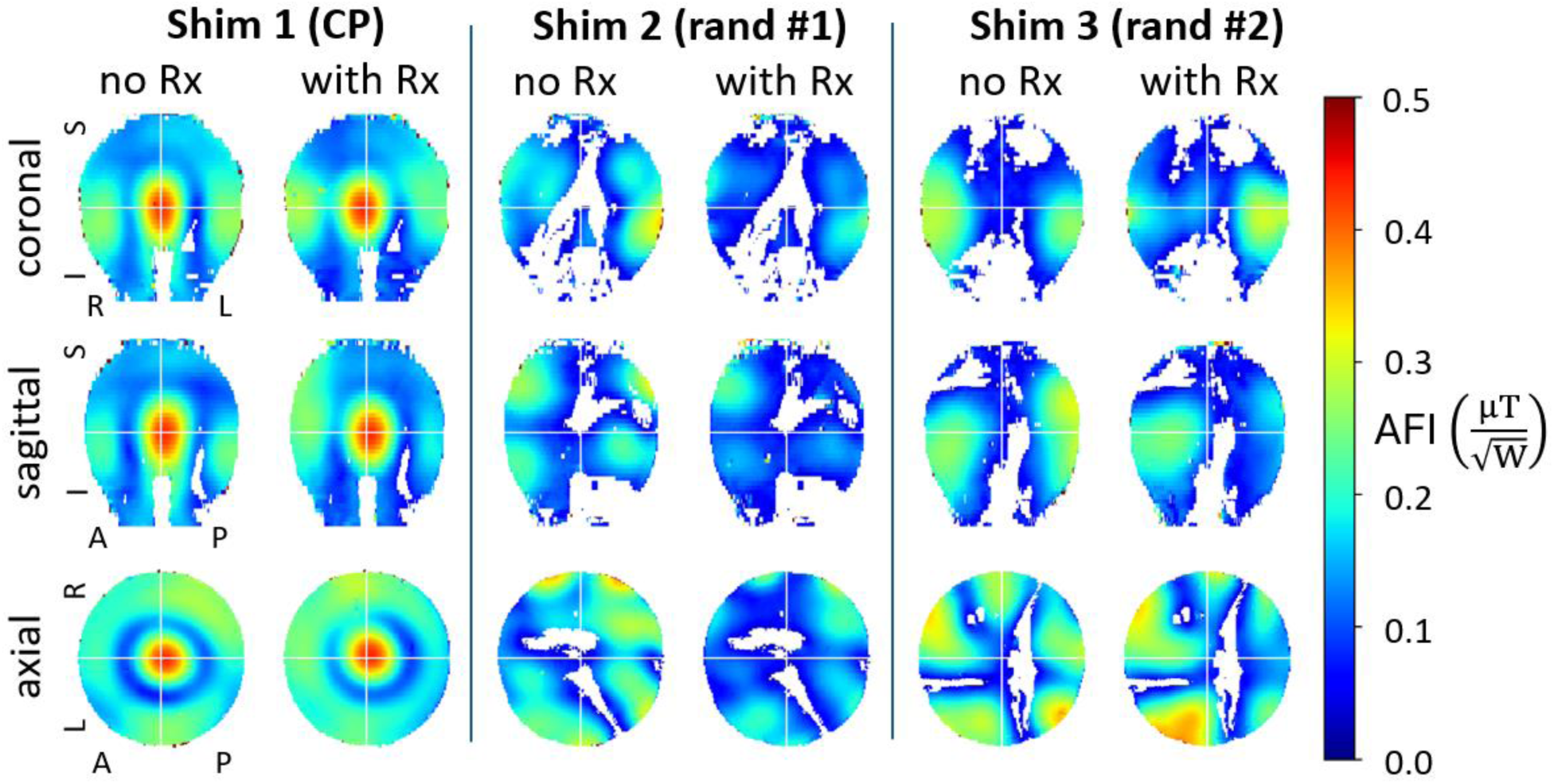
Transmit efficiency 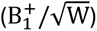 maps with and without the 112Rx receive-only array inserted into the 16Tx/Rx transceiver, for three different shim solutions as labelled: CP-like, random #1, and random #2. For all maps, but particularly prominent in the random phase maps, are regions that were excluded from the final map, and appear blank (i.e. white in color); these correspond to regions where B_1_^+^ was too low relative to a preset threshold.

### 3.3 B_1_ Simulation Results and Safety Validation

Experimental and simulated per-channel B***_1_***^+^ maps and the B***_1_***^+^ maps for the CP mode of excitation are presented in Figure 5A and 5B, respectively. The 112Rx insert is not included in the simulation while the experimental data were obtained with the 112Rx inserted.

**Figure 5:**
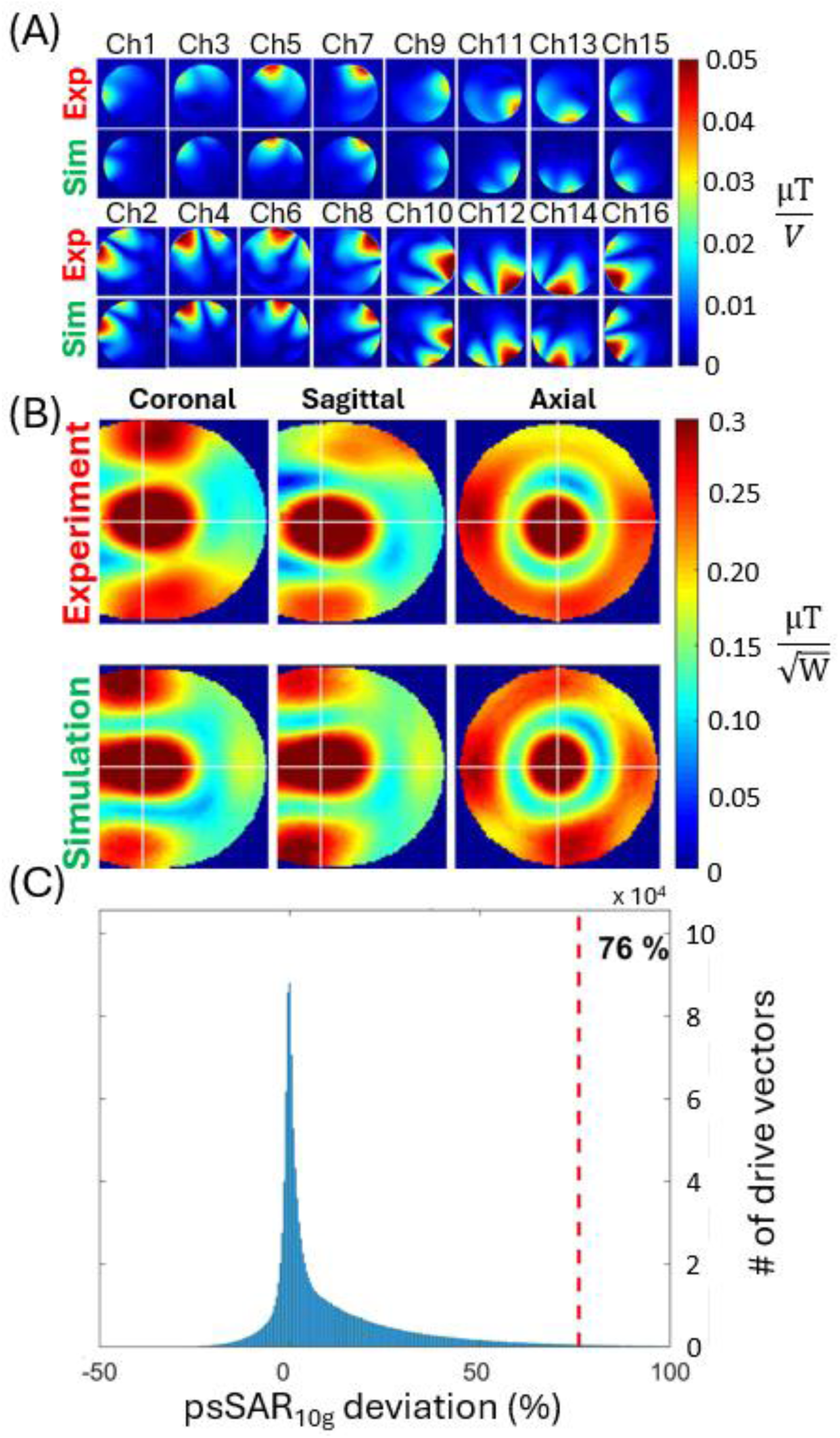
Transmit B_1_ (B_1_^+^) simulation versus experimental data. EM simulation results from Ansys HFSS compared to experimental data for each individual channel **(A)** and for the circularly polarized (CP)-like mode **(B)**. In panel (A) the odd versus even numbered channels correspond to the two rows of azimuthally distributed 8 elements in the 16Tx/Rx array, with even numbers designating the bottom row (towards the feet). Panel **(C)** displays a histogram of the psSAR10g deviation from the psSAR10g of the CP-like mode due to the EM modeling uncertainty. The plot was obtained from Monte-Carlo simulations and shows the distribution of psSAR10g deviation calculated from different RF excitation scenarios (i.e., perturbed around the CP-like mode), satisfying the condition B_1_^+^ NRMSE ≤ 33%. For 99.9% of all RF excitation scenarios psSAR10g deviated less than 76% from the psSAR10g of the CP-like mode.

For the CP mode excitation pattern (Figure 5B), a 33% normalized root-mean-square error (NRMSE) was calculated between simulated and measured B_1_^+^ maps inside the uniform, lightbulb-shaped phantom. This discrepancy necessitated calculation of the EM modeling error (*e_EMM_* ) for defining safe operational limits in human studies; error between simulated and measured B_1_^+^ maps for the CP mode excitation was propagated to the peak-SAR 10g error (psSAR10g) using 10^6^ Monte-Carlo simulations as detailed in ^64^. Figure 5C illustrates the histogram of the propagated SAR error for the CP mode. The *e_EMM_* for this coil was determined to be 76%, resulting in a safety factor of 1.92, including an inter-subject variability (*e_ISV_* )^65^ of 50% and power monitoring uncertainty ( *e_PM_* ) of 15%. By scaling the VOPs with the safety factor, a total input power limit of ∼20W was calculated for the CP-like excitation in the first-level operating mode, in accordance with IEC guidelines^66^.

### 3.4 SNR and Parallel Imaging Performance

Unaccelerated intrinsic SNR maps obtained in the lightbulb-shaped phantom are shown in Figure 6 for the 10.5T 128Rx coil with comparison to 7T and 10.5T 64Rx arrays^13,25^ and a 10.5T 80Rx array (64Rx+16Tx/Rx)^13^; Figure 6A illustrate SNR maps on three orthogonal slices through the lightbulb-shaped phantom.

**Figure 6:**
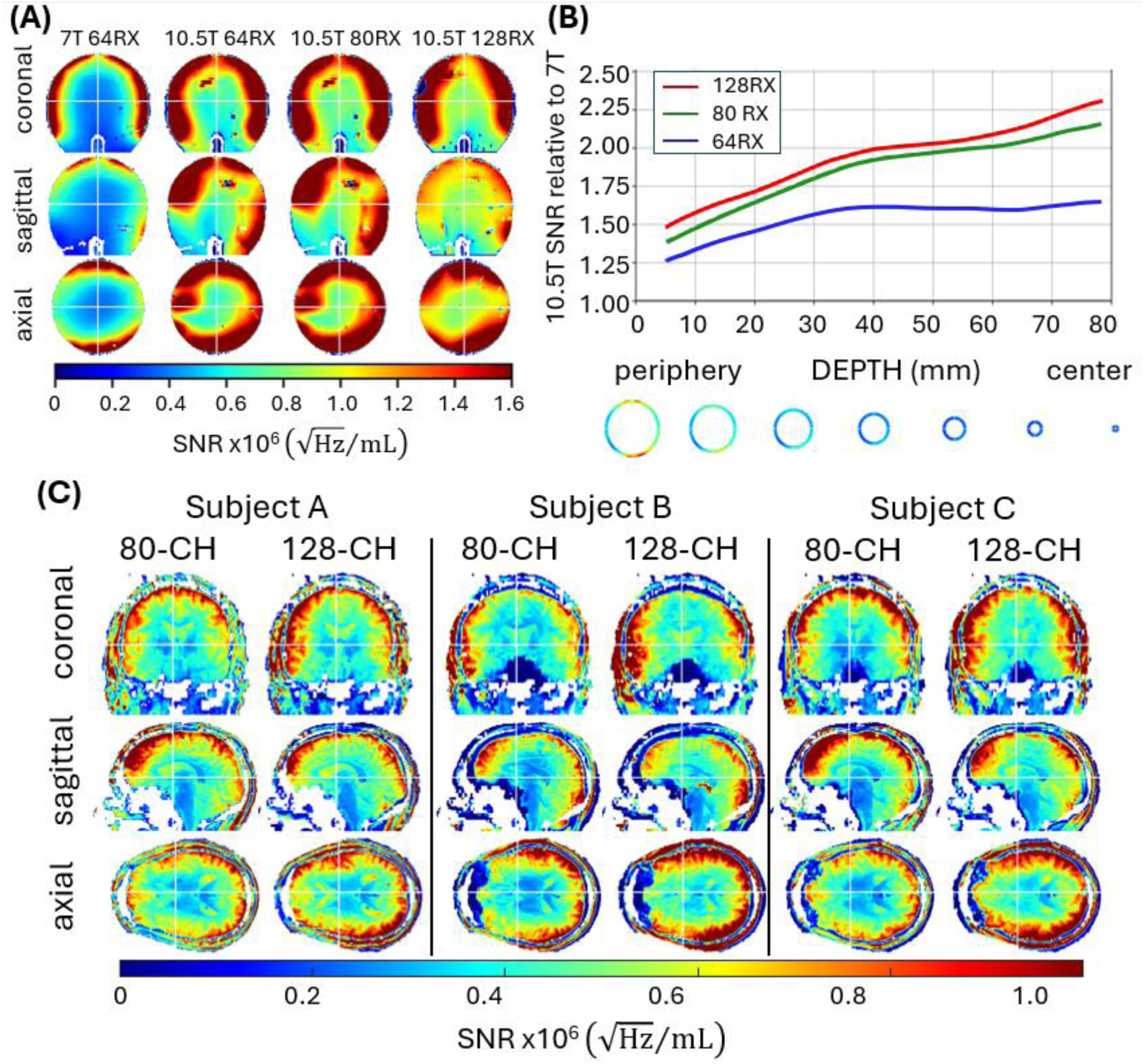
(A) experimental intrinsic SNR maps for a 7T 64Rx, 10.5T 64Rx, 10.5T 80Rx, and the 10.5T 128Rx arrays in three orthogonal slices in the lightbulb-shaped phantom; the 7T and 10,5T 64Rx arrays have identical layouts .(B) the ratio of intrinsic SNR averaged over concentric shells according to shell depth for the 10.5T coils vs the 7T 64Rx coil. The outermost shell was first defined so as to adhere to the outer boundaries of the phantom. This shell was then progressively shrunk. Axial view of the shells are shown below the plot. (C) Intrinsic SNR maps on the same three human volunteers for the 10.5T 80Rx and 128Rx coils.

SNR maps obtained in the phantom were converted to a line plot by using 1 cm thick shells starting at the periphery and subsequently shrinking the shell progressively to obtain concentric shells moving toward the center of the phantom (as illustrated in axial slices below Figure 6B); SNR of each concentric shell was averaged and a ratio taken relative to the SNR obtained with the 7T 64Rx array; this ratio was then plotted as a function of distance from the periphery of the phantom. We previously described this way of presenting SNR data.^13^ The 64Rx arrays at 10.5T and 7T were identical in coil layout. Data for the 7T and 10.5T 64Rx arrays and the 10.5T 80Rx array are reproduced from our previous paper^13,25^ for easy comparison to the 128Rx data.

Compared to the 7T 64Rx array, the 128Rx coil has ∼1.5-fold higher SNR in the periphery and ∼2.5-fold higher SNR centrally. Line plots in Figure 6B also demonstrate that 128Rx has slightly better SNR compared to 80Rx peripherally at 10.5T; in addition, there is a robust ∼15% increase in central SNR with the 128Rx array *versus* the 80Rx array at 10.5T (Figure 6B).

Figure 6C illustrates the SNR for the 128Rx vs the 80Rx arrays at 10.5T in the human head for three different individuals. For each subject, SNR gains in the periphery and in the center of the head are observable for the 128Rx array relative to the 80Rx array, though a slight degradation is observed for peripheral SNR in the frontal areas for the 128Rx versus the 80Rx array.

Parallel imaging performance is demonstrated as 1/g maps in Figure 7. This figure reproduces data we previously published^41^ for 32Rx and 64Rx arrays at 7T for convenience. The remarkable reduction in g-factor noise is observed in going from 7T to 10.5T. At 10.5T, performance of the 128Rx is better in the posterior part of the head near the visual cortex, while being marginally worse in other areas compared to 80Rx.^13,25^

**Figure 7:**
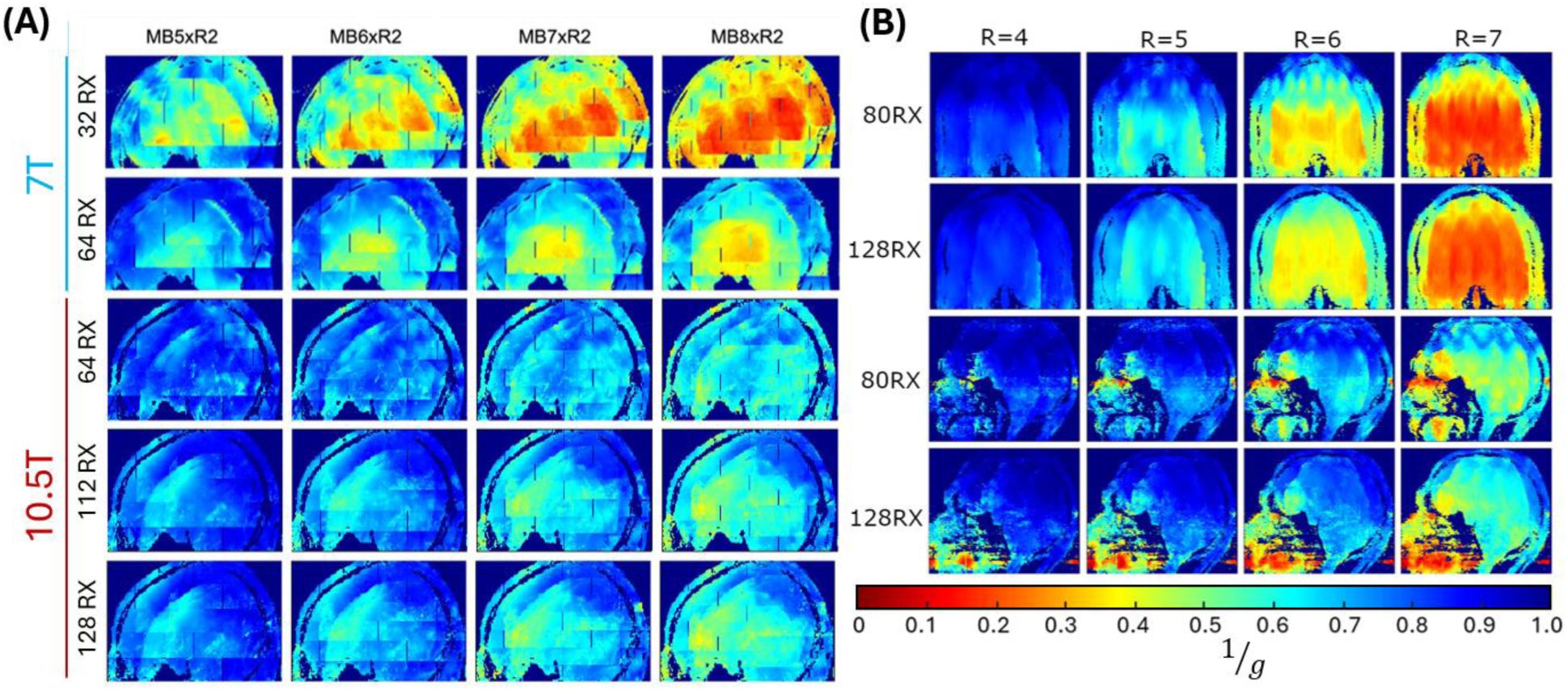
1/g maps, MIP over an 80 mm slab. **(A)** comparing 7T 32Rx and 64Rx arrays (reproduced from Ugurbil et al. 2019, reference 41) to 10.5T 64Rx, 112Rx, and 128Rx arrays for R=2 and increasing MB factors from 5 to 8. The 32Rx array is the industry standard NOVA 32 channel RF coil. **(B)** 10.5T 80Rx and 128Rx comparison for different 1D reduction factors in the left-right direction (coronal plane; top two rows) and anterior-posterior direction (sagittal plane; bottom two rows).

Figure 8 shows the performance of the 128Rx array at 10.5T as a ratio of measured intrinsic SNR of the array (SNR***_array_***) relative to uiSNR in the lightbulb-shaped phantom. The 7T 64Rx and 10.5T 80Rx data are reproduced for convenience from our previous papers.^13,25^ The uiSNR was calculated separately for 7T and 10.5T, since they are not the same. Maps of the SNR***_array_***/uiSNR ratio in a central axial slice are illustrated in Figure 8A. In a 2 cm diameter central sphere (black circles in Figure 8A), the 64Rx arrays with the identical receiver loop layout and proximity to the sample capture a significantly higher fraction of the uiSNR at 7T compared to 10.5T (76% vs 56%)(Figure 8B), in agreement with EM simulations.^25^ Incorporating a 16Tx/Rx array into the same 64Rx-only array (to make an 80Rx array) increases its central SNR performance to 71%.^13^ Adding more channels to the receive only array (112Rx vs 64Rx) leads to higher SNR centrally (Figure 8C) and combining with the same 16Tx/Rx as in the 80Rx array to make a 128Rx array raises the fraction of central uiSNR captured to 77% at 10.5T. Gains in the fractional uiSNR captured in the 128Rx go beyond the 2 cm diameter sphere for which the bar graphs in Figure 8B were calculated (Figure 8A).

**Figure 8:**
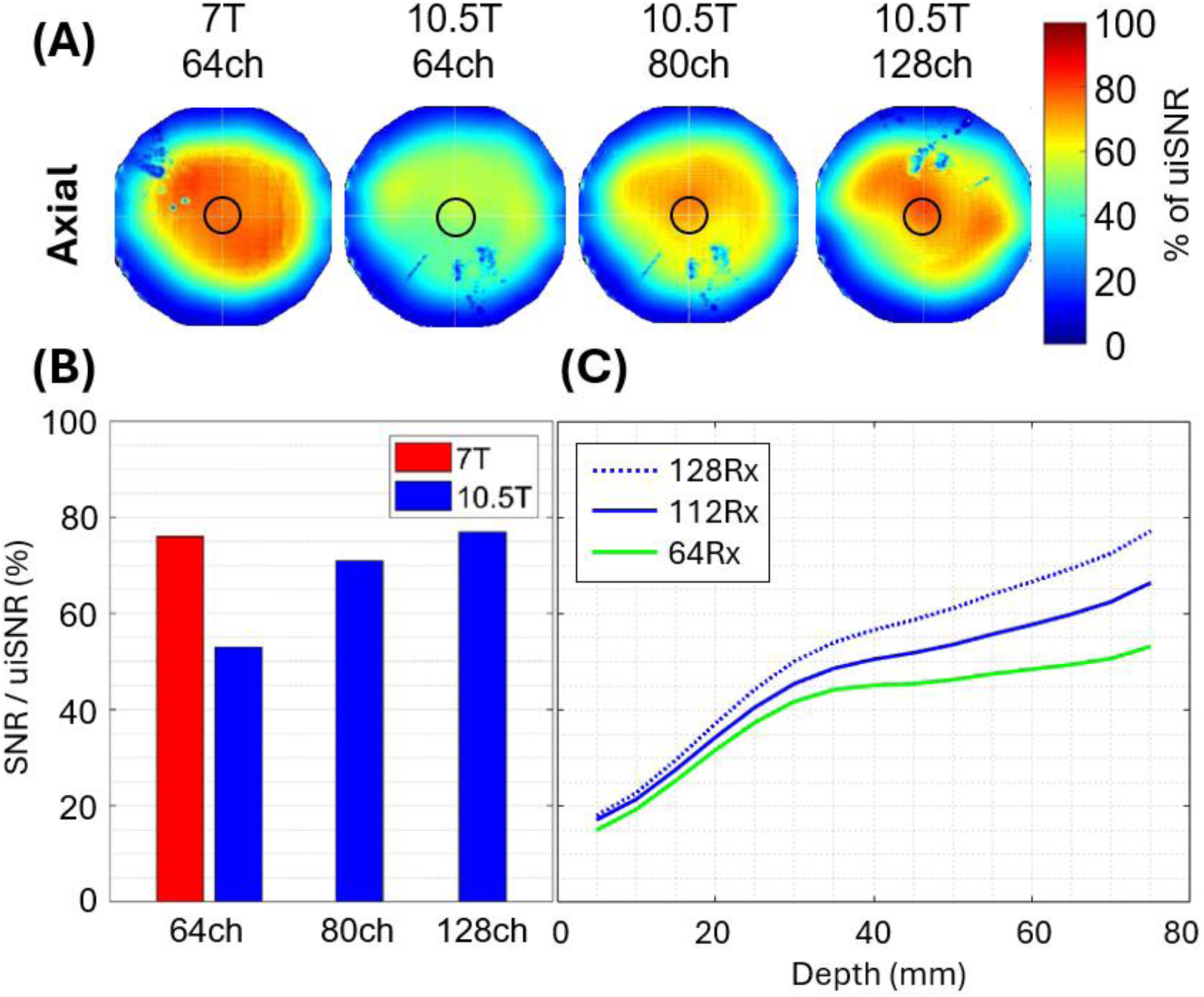
Ratio of the intrinsic experimental SNR of the array relative to uiSNR (SNR***_array_***/uiSNR) in the lightbulb-shaped phantom, illustrating the fraction of the uiSNR captured by the 64Rx arrays at 7T and 10.5T, 80RX array at 10.5T and 128Rx array at 10.5T. **(A)** displays SNR***_array_***/uiSNR ratio in an axial slice as a map. **(B)** shows percent of uiSNR captured by the array as bar-graphs in a central 2 cm diameter spherical volume, the location of which is indicated by the central black circle in the axial maps shown in (A). **(C)** is percent of uiSNR captured (same scale as panel B) vs depth at 10.5T for 64Rx, 112Rx, and the 128Rx arrays.

### 3.5 In vivo Imaging

The in vivo imaging performance of this 128Rx array at 10.5T is showcased in Figures 9 and 10. Figure 9 illustrates high resolution GRE images in a sagittal and axial plane, showing exquisite depiction of the cerebellar structures. See Figure S3 for expanded versions.

**Figure 9:**
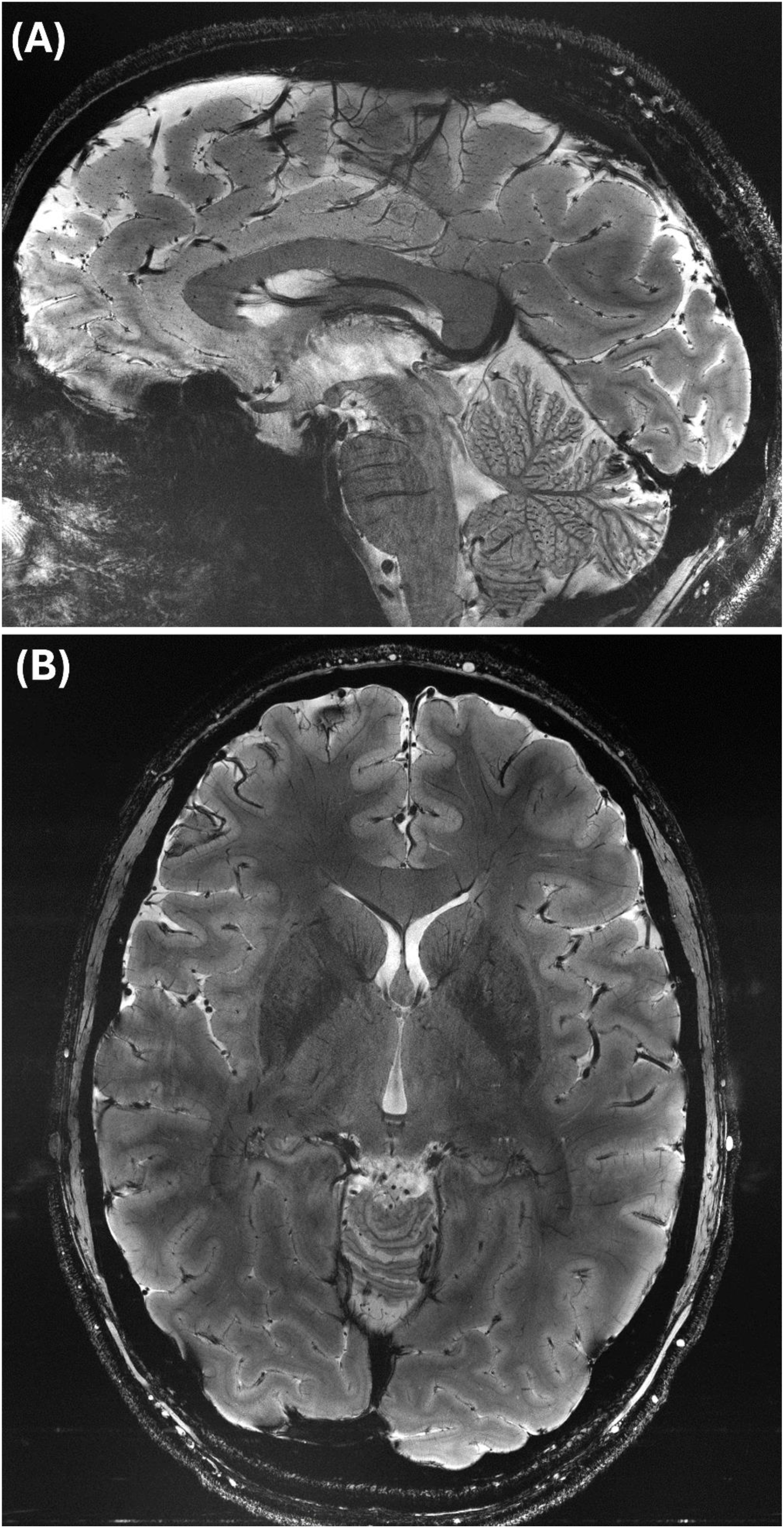
Representative sagittal **(A)** and axial **(B)** slices of a 0.2x0.2 mm^2^ in-plane resolution gradient recalled echo acquisition with a slice thickness of 1 mm, R=2, TA: ∼5 minutes. Note that (A) is cropped slightly to maximize the presence of the brain within the panel.

**Figure 10:**
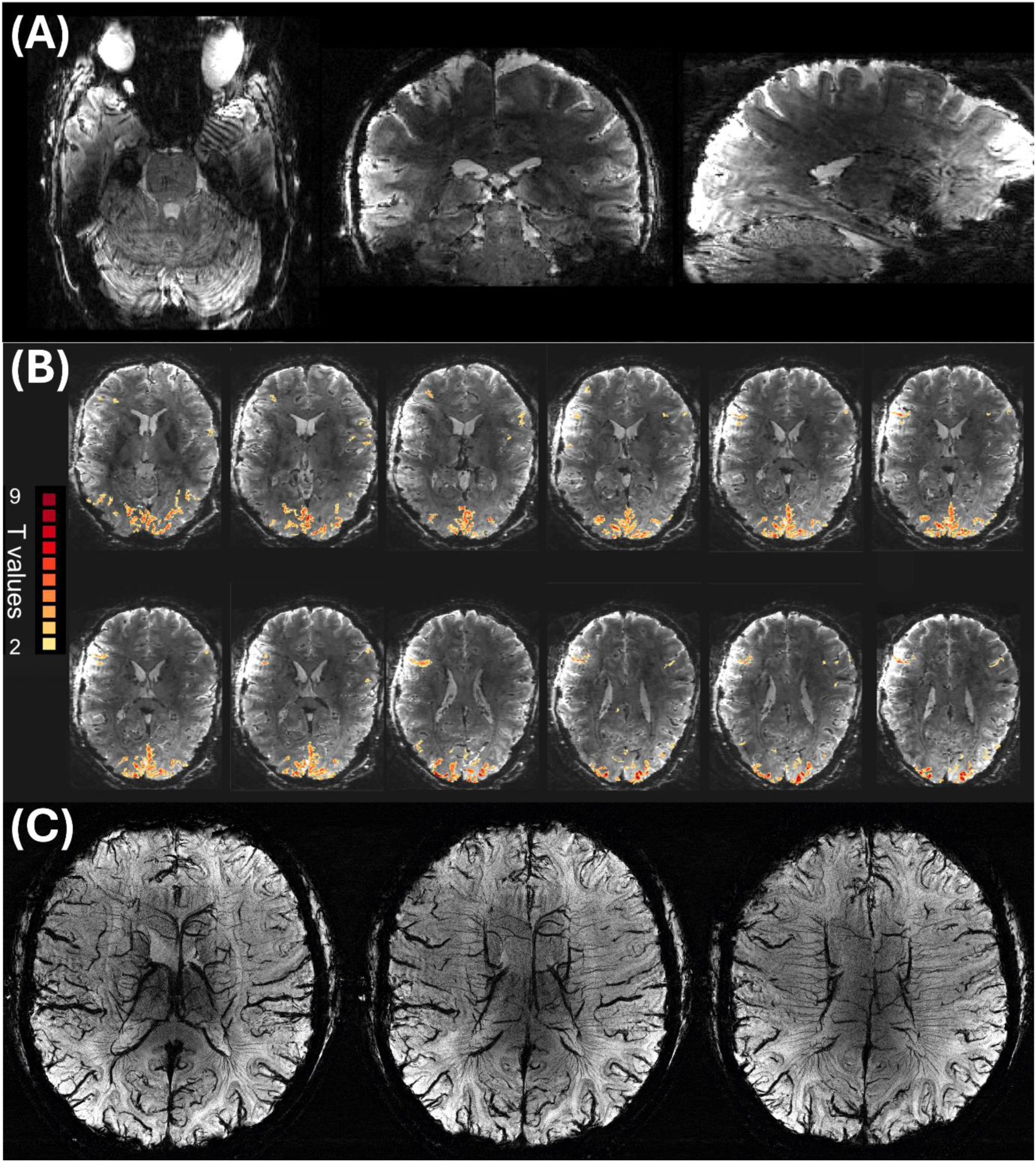
**(A)** Whole-brain 3D GRE EPI images acquired with 0.8 mm isotropic resolution. No denoising or averaging, iPAT 4, FOV: 187 mm x 200 mm x 112 mm (Matrix size 234 x 250 x 144), TE/TR: 20/83 ms, ETL:44, VAT: ∼12 s **(B)** fMRI data obtained using visual stimulation with the same pulse sequence and resolution as (A) but over a 35 mm slab covering the visual cortex. **(C)** SWI images, mIP over a 9.1 mm slab, acquired in 36 contiguous axial-coronal oblique slices with 0.2x0.2 mm^2^ in plane x1.3 mm^3^ slice resolution; FOV: 215 x 188 x 46.8 mm³, TE/TR=18/35 ms, iPAT 5, TA: ∼5 min.

Figure 10A illustrates three orthogonal slices, demonstrating whole head coverage with 3D-GRE EPI with 0.8 mm isotropic resolution at 10.5T obtained with 5-fold in-plane acceleration. Figure 10B shows GRE-EPI BOLD fMRI (visual stimulation) at 0.8 mm resolution obtained with the same 3D sequence but over a slab through the visual cortex. The fMRI data were *not* denoised using algorithms such as NORDIC^67,68^ in order to demonstrate the native fMRI capability of 10.5T. Finally Figure 10C shows mIP images of 0.2x0.2 mm^2^ in-plane resolution susceptibility weighted images (SWI).

## 4. DISCUSSION

In this paper, we describe an effort to optimize human brain imaging at 10.5T by developing a 128Rx/16Tx head array. This is a complex array operating at a very high frequency where no prior experience exists. As such, it is only a preliminary exploration of the challenges encountered in developing such arrays. Initial results for this array were presented in the 2021 and 2023 annual meetings of the ISMRM.^69,70^

### Receive-only array construction

The number of loops in a multi-channel head array cannot be increased *ad infinitum*. As the number of loops distributed over a fixed surface area increases, the loop diameters decrease and ultimately compromises SNR. The point of diminishing return to more channels can be explored using the Q-ratio metric (Figure 2). A loop with a Q-ratio below 2.0 is coil noise dominant, where the electronics contribute more noise than the sample. For optimal SNR performance, the loop should be significantly sample noise dominated with a Q-ratio greatly exceeding 2.0. Fulfilling this condition depends on the resonance frequency, coil diameter, and proximity to the sample, as demonstrated in Figure 2. In general, higher magnetic fields support smaller loops and hence higher number of channels, as shown for 10.5T vs. 7T data in Figure 2B, and previously for 7T vs 3T.^45^ Distance to the sample, however, also plays a prominent role (Figure 2). This is particularly relevant since elements of an array designed for human head imaging are generally not equidistant to the surface of head.

For the ∼4 cm diameter loops employed in the inner 112Rx array of the 128Rx/16Tx coil, Q-ratios when examined as a simple tuned coil (Figure 2) were greater than 2 even at 50 cm distance from the load. In the coil however, the Q-ratio varied between less than 2 to 3 depending on whether the loops were located at the back of the head or over the forehead. These Q-ratios are lower than optimal due to a low Q***_U_*** of 120, which is a product of both the FPC conductors used as well as detune circuitry parasitics. A comparable silver-plated copper wire loop (e.g. the eye loops) using identical lumped components still suffers significantly from detune circuitry parasitics with Q***_U_***∼160. Thus, while data in Figure 2 indicate general trends and importance of distance to load and the resonance frequency, real implementations on the coil impose additional constraints on the performance.

### Transmitter-Receiver Interactions

In such a complex head coil, the receive-only array could significantly alter the transmit field patterns, particularly at UHFs. At 447 MHz, the free space wavelength is ∼67 cm. Consequently, it is imperative to consider the non-uniform distribution of current in resonant structures and the tendency of conductors at a significant fraction of a wavelength to begin radiating/scattering RF. These interactions are both an impediment to deliver power to the sample and a major obstruction to developing accurate EM models necessary for defining safe operation limits.

Because of the need for accurate EM models of the transmitter, the first human images at 10.5T were of the torso due to the relative simplicity of these transceive coils.^71,72,20,73^ Excellent results were also obtained for a simple 8-channel dipole transceive array for head imaging.^21^ However, initial experiences highlighted the difficulty of achieving this with ensembles composed of separate circumscribing transmitters and inner receive-only arrays. Inability to accurately predict transmit B***_1_*** (B***_1_***^+^) fields from the EM model of the transmitter alone necessitates either incorporating a large safety factor into the power limit calculations or modeling the entire receive array, including all cables, cable traps, etc. together with the transmitter to achieve better accuracy. This would be a daunting task for the 128Rx/16Tx array as can be appreciated from Figure 1. Consequently, several innovations were necessary to reduce interactions of the inner 112Rx array on the transmit field patterns. This involved miniaturizing the footprint of on-coil preamplifier boards, pulling off-coil preamplifier boards entirely out of the transmitter volume, and using cable traps every 𝜆/16 on coaxial cables within the transmitter’s field. The resultant B***_1_***^+^ patterns obtained with and without the 112Rx array in place (Figure 4) were similar, and the B***_1_***^+^ predicted by the EM model of the transceiver was in good agreement with the experimentally measured data (Figure 5). The residual discrepancy was dealt with using the approach described in the Supporting Information. Consequently, the coil was approved for human imaging, enabling the demonstration of excellent imaging results (Figures 9 and 10). This included T***_1_***-weighted images obtained with MP2RAGE which requires high peak-power 180° pulses (Figure S4). This is in contrast to the performance of the recent 7T 128Rx coil;^45^ it was reported that “the 128-channel receive array hardware caused about a 33.6% loss in efficiency at the center of the FOV and also introduced a crescent-shaped region of low transmit efficiency” (see Figure S4 in ^45^).

### SNR

Our recent EM modeling work and accompanying experimental data demonstrated that 32Rx or 64Rx arrays composed of loops capture a much smaller fraction of the central uiSNR at 10.5T compared to 7T.^25^ We also showed that an 80Rx array, composed of a 64Rx-only inner array and a second layer of 16Tx/Rx elements led to significant increases in central SNR relative to the 64Rx array alone and brought the fraction of central uiSNR captured at 10.5T close to levels achieved by a 7T 64Rx array of identical construction.^13^ Here we demonstrate that at 10.5T, the 112Rx receive-only array relative to a 64Rx array also yields SNR gains centrally as well as peripherally (Figure 8C). Counterintuitive from experiences derived from <7T, the largest gains are centrally located. Using the same strategy as in the 80Rx array to engage the transmitter loops as receivers provides another incremental gain. Despite the use of an identical transceiver, the gains in central uiSNR provided by the transceiver in the 128Rx/16Tx array is not as large as those observed with the 80Rx array both in relative and absolute terms. This suggests a plateau to the maximum fraction of central uiSNR of about 80% for the current constellation of coil design and device noise figures for both 10.5T and 7T (Figure 8).

Of the 10.5T coils examined so far, the 128Rx array yields the highest value for the fraction of central uiSNR captured, approximately matching what the 64Rx array is able to do at 7T (Figure 8). The data in Figure 6 thus can be taken as a measure of the intrinsic field-dependent SNR gains centrally in going from 7 to 10.5T for a phantom approximating the human head. This yields ∼2.5-fold gain at 10.5T relative to 7T for the center (Figure 6). Within a model where 𝑆𝑁𝑅 ∝ 𝐵_0_^𝑛^, this corresponds to 𝑛 = 2.26. This is in excellent agreement with uiSNR gains at 10.5T relative to 7T calculated in the human head and the same phantom^25^ and also with data derived from 64Rx and 80Rx arrays using corrections based on the performance of each coil relative to uiSNR.^13,25^ The field-dependent central SNR increase measured for the 128Rx array is also in good agreement with results reported for the center of a sphere^23^ derived from data obtained with birdcage coils.

Compared to the 80Rx array, the 128Rx coil in general provides robust SNR gains both in the center and the periphery. However, the gains in the periphery are not uniform in space. This is particularly evident in the human data (Figure 6). However, the 128Rx and 80Rx arrays have several differences that make a direct comparison difficult. Most significant is that the formers used for inner arrays are not identical and the 128Rx array former is larger particularly in the superior region (Figure S1). For practical reasons, the majority of the loop conductors for the 128Rx array were constructed from FPCs. Q-factors indicate that coil noise is increased for the FPC coils, consistent with previously reported data.^38,74^ Additionally, while coil diameter generally decreases with increasing channel count, this does not hold true if coil layout changes. The 128Rx array uses round loops with overlap in all directions while the 80RX array uses rectangular loops with only partial overlapping. The 128Rx array uses a standard round loop layout with overlap decoupling to all nearest neighbor loops with no mitigation to next-nearest neighbor coupling, instead relying upon preamp decoupling. The 80Rx coil, by contrast, has lower next-nearest neighbor coupling since loops are not overlapped in the azimuthal direction; however nearest neighbors rely solely upon preamp decoupling. Even preamplifier decoupling is approached differently. Because of Tx/Rx interaction mitigation strategies employed and the presence of on- coil preamps, the 128Rx array relies upon more lumped-component phase shifting networks than the 80Rx array. These networks have potential for introducing additional losses if not optimally tuned.

All these differences are contributing factors to the lower-than-expected peripheral SNR gains for the higher channel count array. However, as a brain imaging coil, the 128Rx array still achieves higher central SNR which yields excellent practical imaging performance.

### Parallel Imaging

Parallel imaging performance at 10.5T was exceptional compared to a 7T (Figure 7). However, it was only marginally better for the 128Rx array compared to the 80Rx array at 10.5T for the orientations examined (Figure 7). As discussed, however, there are numerous differences between the 80Rx and 128Rx arrays that account for this including former dimensions and loop layout. It is likely that non-overlapped loops in the 80Rx produce more complex and distinct sensitivity profiles than fully overlapped loops in the 128Rx. These differences notwithstanding, this finding is in close agreement with previously published results at 7T showing a lack of major differences in g-factor in going from 64Rx to 96Rx arrays.^75^

## 5. CONCLUSIONS

We present a 128Rx/16Tx array for 10.5T head imaging as an important next step in the effort to fully realizing the SNR advantage available by going to higher magnetic fields. The 128Rx array to date provides the best performance in terms of capturing the central uiSNR available at 10.5T, delivering the field dependent gains expected from uiSNR calculations. We successfully demonstrate that the complexity of building such a dense array within a circumscribing transmitter was successfully managed which is a key technical achievement that will likely see adoption by future UHF receive arrays. While certain design decisions resulted in lower-than-expected improvements in peripheral SNR compared to our 80Rx 10.5T coil, this cannot be generalized to high channel count coils and is specific to this array.

## Supporting information

3D CAD

## ACKNOWLEDGEMENTS

The authors would like to thank Dr. Shajan Gunamony for his decades of support, kindness and collaboration, particularly for productive discussions surrounding preamp specifications as well as mechanical and electrical design of our transmit arrays.

Our summer interns Erne Habeggar McCabe and Marybelle Kim proved to be talented technicians and were responsible for replication of the large quantity of cable traps, preamp boards, and feed boards needed in this coil.

We would like to honor and remember Pierre-Francois Van de Moortele for his invaluable contributions to the field in general including to 10.5T which paved the way for this work.

## SUPPORTING INFORMATION FIGURE CAPTIONS

Figure S1: Differences between the 80Rx and 128Rx coil former (64Rx-only insert vs 112Rx- only insert, respectively). The top two images show a cross-section of the former along the sagittal plane with coronal and trans-axial shown on the bottom. The 80Rx former is shown in red while 128RX is depicted by blue (and gray for the top-left figure).

Figure S2: Photos of the unpopulated 112Rx array with a photo of the corresponding flexible printed circuits unwrapped below and the associated loop layout at the very bottom.

Figure S3A: Expanded version of Figure 9A, showing an axial slice from a 0.2x0.2 mm^2^ in-plane resolution gradient recalled echo acquisition with a slice thickness of 1 mm, R=2, TA: ∼5 minutes. The image is cropped slightly to maximize the presence of the brain within the panel.

Figure S3B: Expanded version of Figure 9B, showing an axial slice from a 0.2x0.2 mm^2^ in-plane resolution gradient recalled echo acquisition with a slice thickness of 1 mm, R=2, TA: ∼5 minutes.

Figure S4: 0.8mm isotropic resolution MP2RAGE images of two separate human subjects (A and B). TE/TR = 1.89/5000 ms, TA: 6 min 35 sec, FOV: 240 x 256 x 153.6 mm³ (Matrix size: 300 x 320 x 192)

