## Supporting Figures and Information for "A 128-channel receive array with enhanced SNR performance for 10.5 tesla brain imaging"

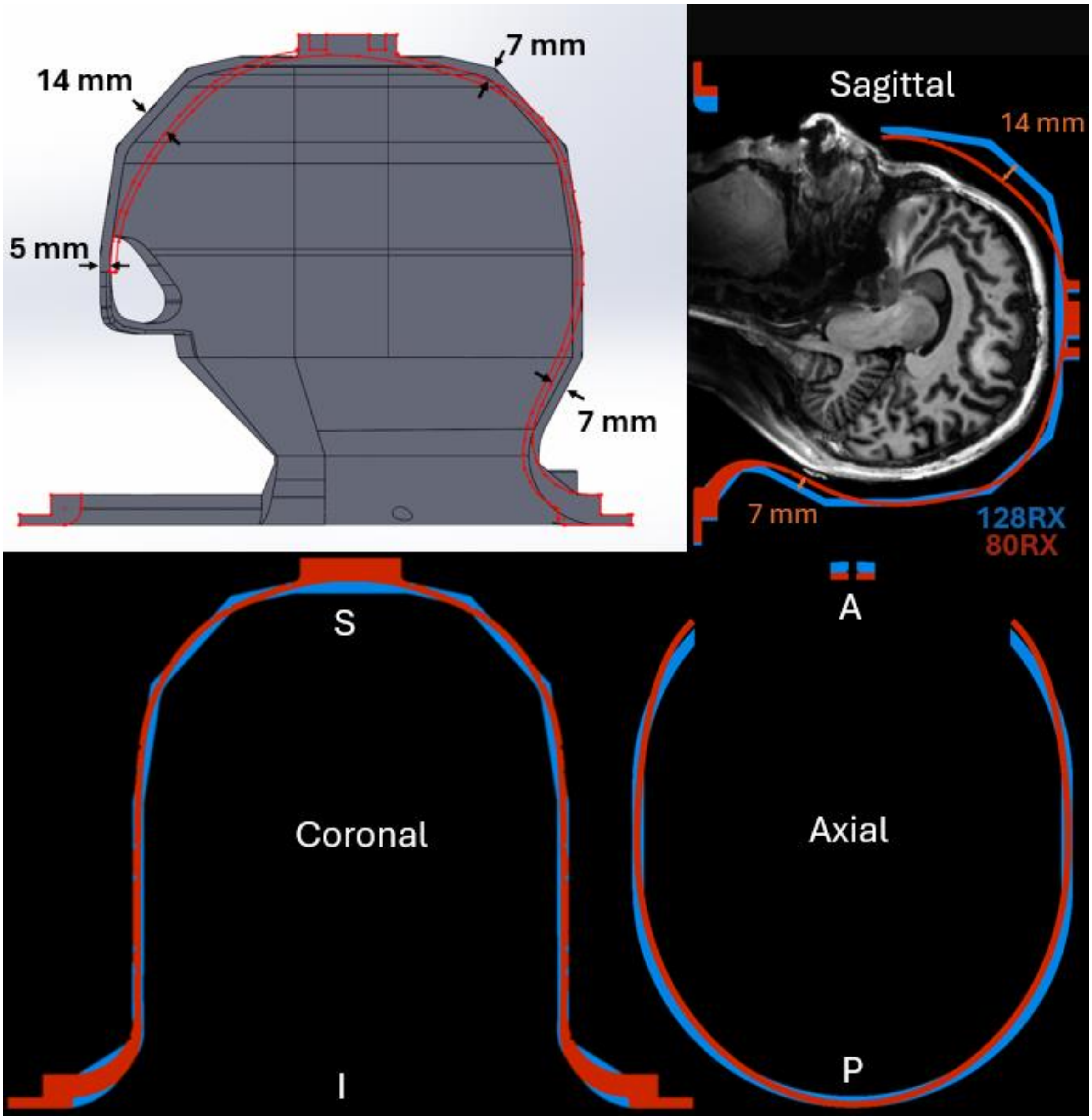


**Figure S1:** Differences between the 80Rx and 128Rx coil former (64Rx-only insert vs 112Rx-only insert, respectively). The top two images show a cross-section of the former along the sagittal plane with coronal and trans-axial shown on the bottom. The 80Rx former is shown in red while 128RX is depicted by blue (and gray for the top-left figure).

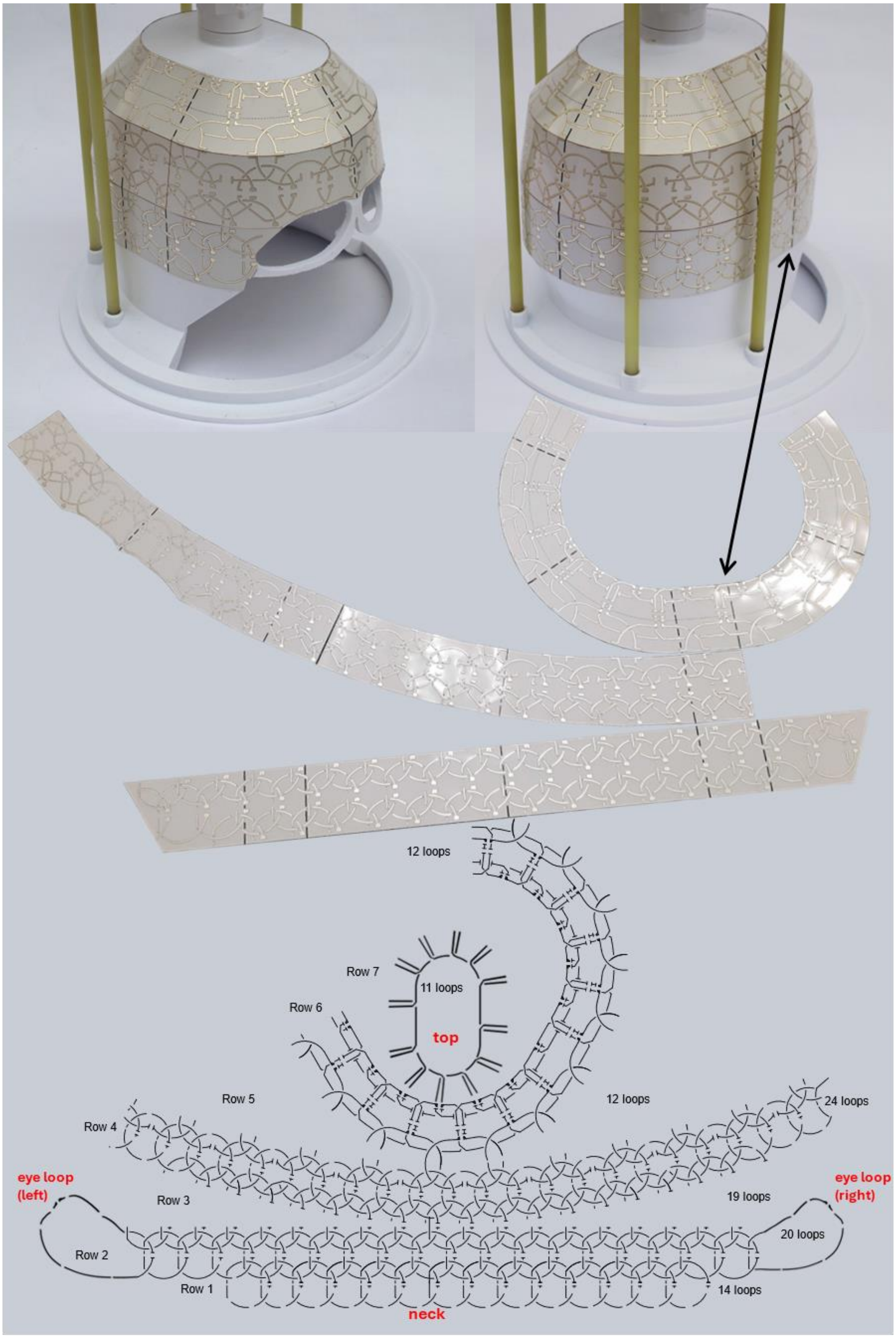


**Figure S2:** Photos of the unpopulated 112Rx array with a photo of the corresponding flexible printed circuits unwrapped below and the associated loop layout at the very bottom.

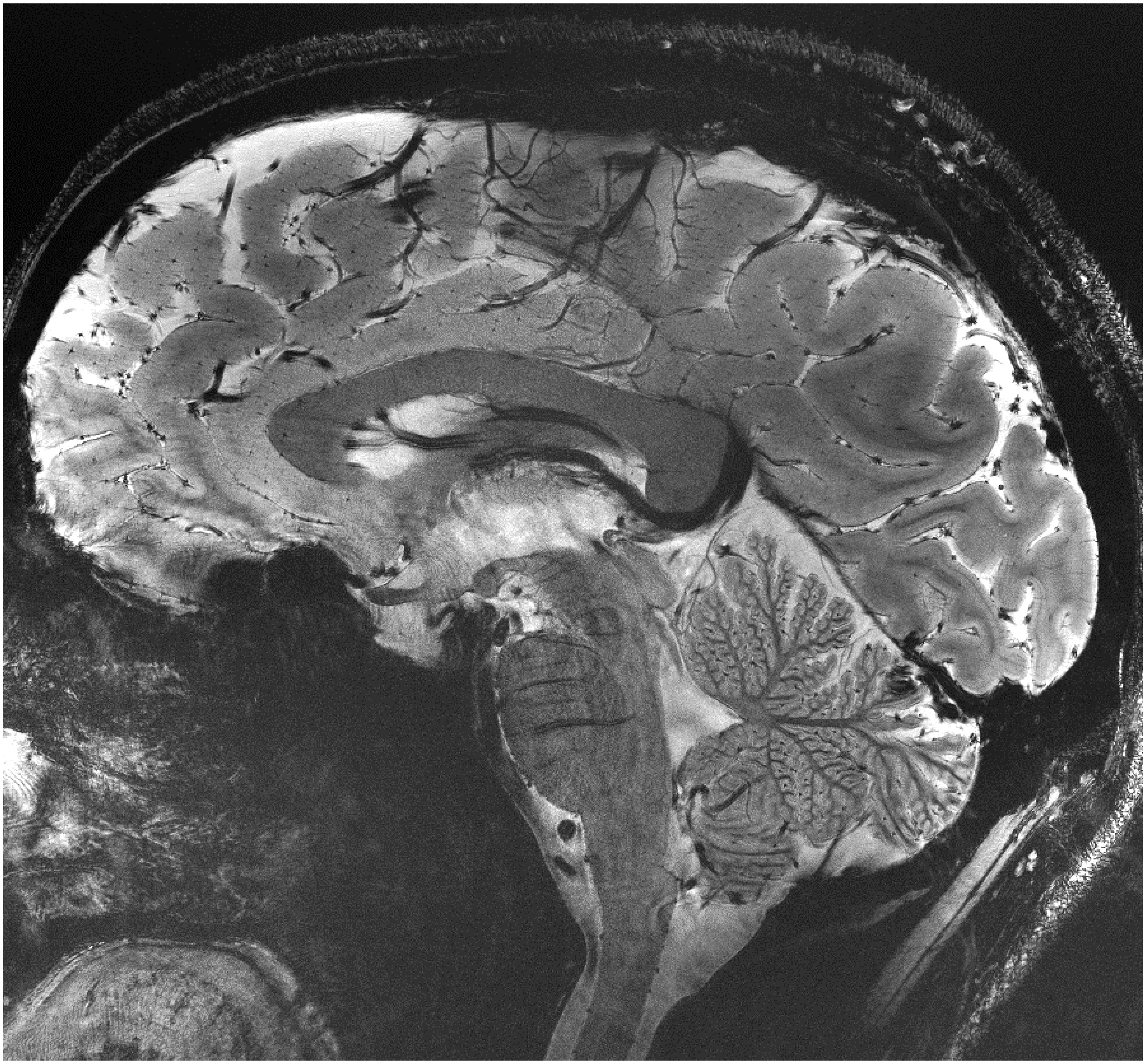

**Figure S3A**: Expanded version of Figure 9A, showing an axial slice from a 0.2x0.2 $mm^2$ in-plane resolution gradient recalled echo acquisition with a slice thickness of 1 mm, R=2, TA: ~5 minutes. The image is cropped slightly to maximize the presence of the brain within the panel.

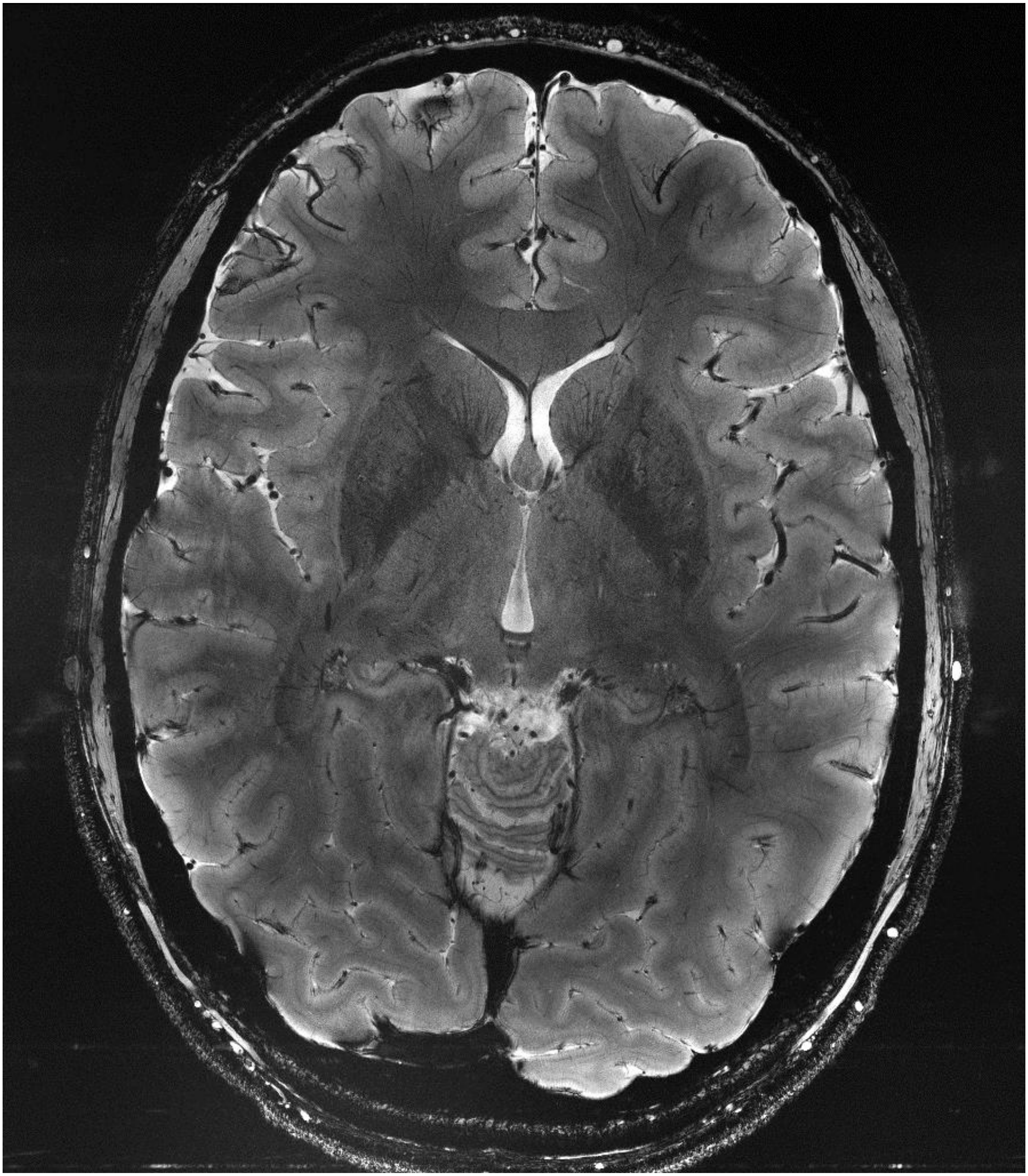

**Figure S3B**: Expanded version of Figure 9B, showing an axial slice from a 0.2x0.2 $mm^2$ in-plane resolution gradient recalled echo acquisition with a slice thickness of 1 mm, R=2, TA: ~5 minutes.

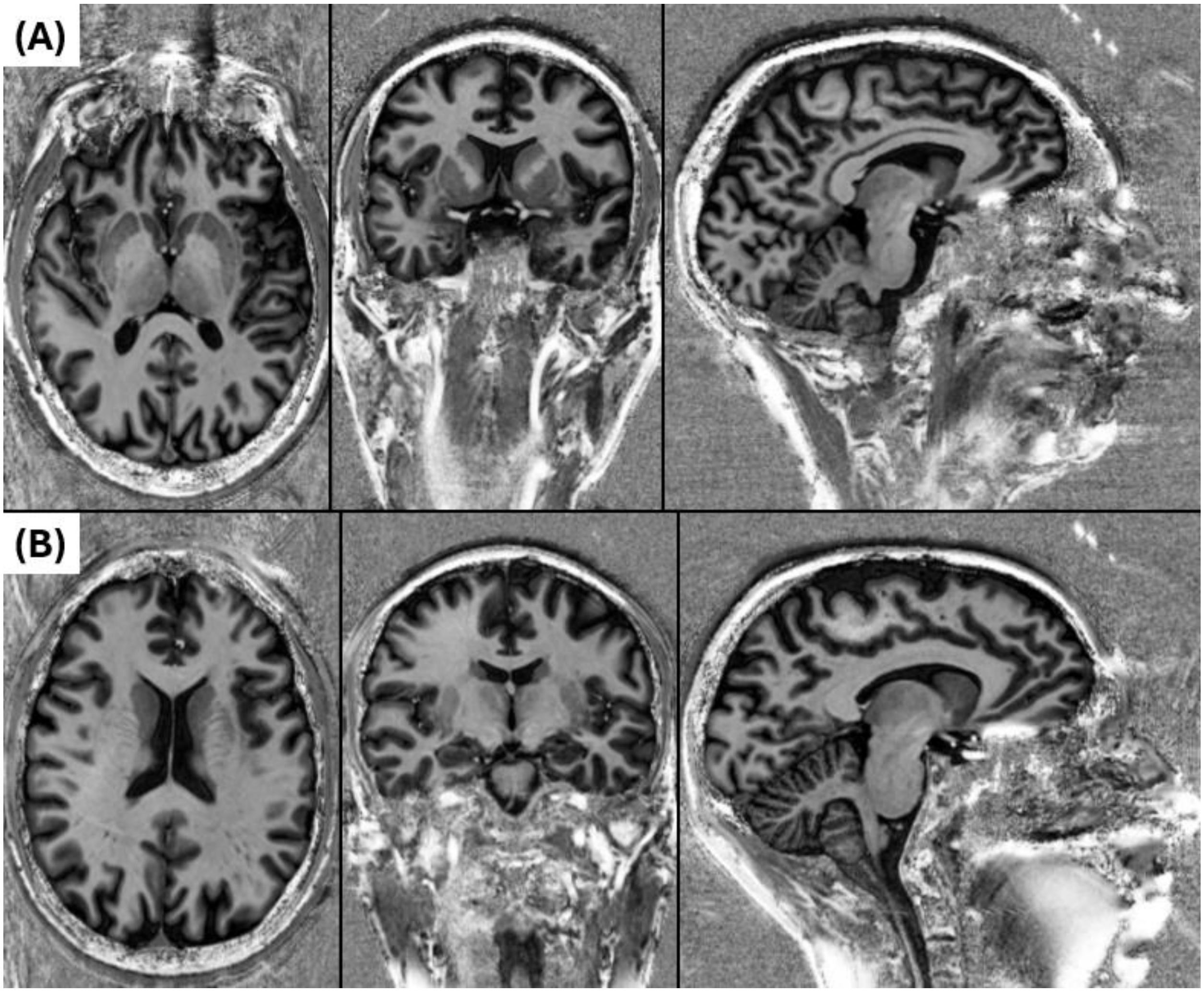


**Figure S4:** 0.8mm isotropic resolution MP2RAGE images of two separate human subjects (A and B). TE/TR = 1.89/5000 ms, TA: 6 min 35 sec, FOV: 240 x 256 x 153.6 mm³ (Matrix size: 300 x 320 x 192)

### *2.1.2 Transceiver Array*

RF transmission for spin excitation was accomplished with a 16-channel transceiver (16Tx/Rx) array set up to transmit and receive with each of its elements. This array is based on self-decoupled (SD) loops,[1] and is identical to the transceiver previously described[2,3]. The coil was constructed onto a 3D printed PETG plastic former (ID x OD x L = 27.5 cm x 32 cm x 36 cm). Integrated T/R switches allow the array to be used for both transmit and receive. Two 8-channel system plugs (ODU, Mühldorf, DE) supply the transmit connections to the patient table while a pair of 8-channel receive TIM plugs (Total Imaging Matrix, Siemens, Erlangen, DE) can either plug into the patient table for a simple 16 T/R setup, or plug into sockets on the receive array top plate to routed receive signals through the receive array plugs. Loop conductors were laid out on FPCs which are adhered to the coil former using double-coated film tape (9589, 3M, Maplewood, MN, USA). Loops were laid out in two rows of 8 channels with a 22.5° offset azimuthally between elements in each row (Figure 1B).

### *2.1.3 EM Simulation and Safety Validation*

The CMRR's 10.5T scanner operates under an Investigational Device Exemption (IDE) from the FDA, requiring each RF coil to receive FDA approval before use with humans. A key part of this process is validating the EM models developed for the arrays, which is accomplished using phantoms. In our case, we used a lightbulb-shaped phantom designed to mimic the shape and average electrical properties of the human head and neck. The characteristics of this phantom including its dimensions, internal material composition, and electrical properties was described previously[3,2,4] and the phantom was employed in our previous work for head arrays at 10.5T [3,2,4]. Briefly, the phantom is filled with a composite of polyvinylpyrrolidone (PVP) and saline solution, with electrical properties falling close to an average of several brain tissues at 447 MHz.[5] Dielectric properties were characterized with a DAKS-12 dielectric probe (SPEAG, Zürich, CH). The measured conductivity was 0.65 S/m and the relative permittivity was 47.2.

The RF-related patient safety evaluation of the 128Rx/16Tx array followed a 3-phase workflow.[6,7] The specifics of this workflow for a 10.5T multichannel transmit and receive array was described in detail previously[2,3]; the same procedure and methods were employed in this work. In the first phase, we developed an EM model of the 16Tx/Rx array in HFSS environment (Ansys, Canonsburg, PA, USA) ensuring good agreement between the experimental and simulated $B_1^+$ maps and S-parameters with the coil loaded with the lightbulb-shaped phantom. The second phase involved quantifying the discrepancies between simulated results and experimental $B_1^+$ data and incorporating these into a safety factor. In the final phase, this safety factor was used to scale the 10g-averaged spatially specific absorption rate (Q-matrices) derived from EM simulations with a human head model. During *in vivo* imaging, these Q-matrices were used to calculate the peak 10g-averaged spatial specific absorption rate (psSAR10g) for any particular RF excitation scenario, determining safe power limits in accordance with IEC guidelines.[8]

*Supporting Information Bibliography:*
